# Regulation of early growth response-1 (Egr-1) gene expression by Stat1-independent type I interferon signaling and respiratory viruses

**DOI:** 10.1101/2020.08.14.244897

**Authors:** Chilakamarti V. Ramana

**Affiliations:** Department of Medicine, Dartmouth-Hitchcock Medical Center, Lebanon, NH 03766, USA

**Keywords:** Interferon-*α*/*β*, TNF-*α*, Innate immunity, Stat1-independent pathway, Early Growth Response-1. Influenza viruses, coronaviruses, COVID-19

## Abstract

Respiratory virus infection is one of the leading causes of death in the world. Activation of the Jak-Stat pathway by Interferon-alpha/beta (IFN-*α*/*β*) in lung epithelial cells is critical for innate immunity to respiratory viruses. Genetic and biochemical studies have shown that transcriptional regulation by IFN-*α*/*β* required the formation of Interferon-stimulated gene factor-3 (ISGF-3) complex consisting of Stat1, Stat2, and Irf9 transcription factors. Furthermore, IFN *α*/*β* receptor activates multiple signal transduction pathways in parallel to the Jak-Stat pathway and induces several transcription factors at mRNA levels resulting in the secondary and tertiary rounds of transcription. Transcriptional factor profiling in the transcriptome and RNA analysis revealed that Early growth response-1 (Egr-1) was rapidly induced by IFN-*α*/*β* and Toll-like receptor (TLR) ligands in multiple cell types. Studies in mutant cell lines lacking components of the ISGF-3 complex revealed that IFN-*β* induction of Egr-1 was independent of Stat1, Stat2, or Irf9. Activation of the Mek/Erk-1/2 pathway was implicated in the rapid induction of Egr-1 by IFN-*β* in serum-starved mouse lung epithelial cells. Interrogation of multiple microarray datasets revealed that respiratory viruses including coronaviruses regulated Egr-1 expression in human lung cell lines. Furthermore, Egr-1 inducible genes including transcription factors, mediators of cell growth, and chemokines were differentially regulated in the human lung cell lines after coronavirus infection, and in the lung biopsies of COVID-19 patients. Rapid induction by interferons, TLR ligands, and respiratory viruses suggests a critical role for Egr-1 in antiviral response and inflammation with potential implications for human health and disease.

## 1 Introduction

Interferons (IFNs) are pleiotropic cytokines that play a central role in innate and adaptive immunity (1,2). There are 3 major types of interferons. The type I interferons consists of IFN-alpha/beta (IFN-*α*/*β*). type II interferon represented by interferon-gamma (IFN-γ), and type III interferons represented by interferon lambda (IFN-λ). The biological effects of IFN are mediated mainly by the rapid and dramatic changes in gene expression (3)). Type I IFN signaling involves the binding of IFN-*α*/*β* to its receptor (IFNAR) and activation of receptor-associated Janus protein tyrosine kinases Jak1 and Tyk2 and the phosphorylation of Stat1 and Stat2 to form a heterodimer. This heterodimer associate with the Interferon responsive factor 9 (Irf9) to form the interferon-stimulated gene factor-3 (ISGF-3) complex on interferon-stimulated response elements (ISRE) in the promoters of type I IFN responsive genes to regulate transcription (1,2,4). Stat1 homodimer or heterodimer of Stat1/Stat2 can also bind to Gamma activated sequence (GAS) in the promoter and regulate transcription of some type I IFN responsive genes (1,4). In addition to this classical canonical Jak-Stat pathway, non-canonical pathways involving Stat2, Irf9, and unphosphorylated ISGF-3 components in the regulation of gene expression have been described (5). Furthermore, several signal transduction pathways are activated by IFN receptors in parallel to Jak-Stat including extracellular signal-regulated kinases (Erk-1/2) and phosphoinositide 3’-kinase/Akt pathways (6,7). The first wave of Interferon signaling is followed by induction of transcription factors such as Interferon regulatory factors (IRFs) that sustain the secondary and tertiary transcriptional responses. TLR recognition of pathogens by immune cells results in the production of multiple cytokines such as IFN-*α*/*β*, Tumor Necrosis Factor-*α* (TNF-*α*), and Interleukin-*β* (IL-1*β*) in innate immunity (8). Activation of multiple signal transduction pathways and cross-talk between the pathways enables fine-tuning of gene expression in innate immunity (9). Cross-talk between TNF-*α* and IFN signaling pathways regulate inflammatory gene expression to influence the immune responses (10,11). Clinical significance of the balance between immune modulation and inflammation in signal transduction pathways has been demonstrated in a variety of autoimmune diseases (12,13).

Early growth response 1 (human EGR1/ mouse Egr1; referred in this manuscript as Egr-1) belongs to a family of immediate-early response genes that contain a conserved zinc finger DNA-binding domain and binds to a GC-rich sequence in the promoters of target genes (14). A variety of signals, including serum, growth factors, cytokines, and hormones stimulate Egr-1 expression (15,16). Egr-1 has been shown to play an important role by regulating inflammatory gene expression in a variety of lung diseases and in mouse lung injury models including asthma, emphysema, airway inflammation, and pulmonary fibrosis (17-19). Ischemia-mediated activation of Egr-1 triggers the expression of pivotal regulators of inflammation including the chemokine, adhesion receptor, and pro-coagulant gene expression (20). Egr-1 stimulates chemokine production in interleukin-13 mediated airway inflammation, and remodeling in the lung (21). High levels of expression of Egr-1 and Egr-1 inducible genes were reported in atherosclerosis, an inflammatory disease (22). Egr-1 may have a potential role in liver injury and in acute pancreatitis (23-25). Lipopolysaccharide (LPS) induction of Egr-1 was mediated by the activation of Erk-1/2 pathway and serum response elements (26). In this study, transcription factor profiling in interferon-mediated gene expression data sets and RT-PCR revealed that Egr-1 was rapidly induced by IFN-*α*/*β* and TLR ligands in multiple cell types. Studies in mouse and human fibroblast mutant cell lines revealed that Egr-1 induction by type I interferons was independent of transcription factors Stat1, Stat2 or Irf9. Furthermore, the regulation of Egr-1 by IFN-*β* was mediated by the activation of the Erk-1/2 pathway in serum-starved mouse lung epithelial cells. Respiratory pathogens including coronaviruses (SARS-CoV-1 and 2) and influenza viruses regulated the expression of Egr-1 in human lung cell lines and in lung biopsies and peripheral blood cells of COVID-19 patients, These studies suggest that the regulation of Egr-1 may play an important role in the antiviral response and inflammatory disease.

## 2 Materials and Methods

### Gene expression datasets

Gene expression in response to Interferon, TNF-*α*. and TLR agonist treatment in human peripheral blood mononuclear cells (PBMC), human hepatoma cells (Huh-7), mouse bone marrow-derived macrophages (BMDM) were reported previously (27-29). Supplementary data was downloaded from the Journal publisher websites and Geo datasets were analyzed with the GeoR2R method (NCBI). Cluster analysis was performed using gene expression software tools at www.heatmapper.ca. Gene expression datasets representing human lung cell lines infected with respiratory viruses and from COVID-19 patients were reported previously (30,31). Gene expression resources from Immgen RNA seq SKYLINE were used (http://rstats.immgen.org/Skyline_COVID-19/skyline.html). Outliers of expression were not included in the analysis.

### Cell culture and cytokine treatment

Mouse lung epithelial (MLE-Kd) and macrophage (RAW264.7) cell lines were used (11,32). Human fibrosarcoma cell line (2fTGH) and mutant cell lines lacking Stat1 (U3A), Stat2 (U6A), and IRF9 (U2A) were described previously (33). Wild -type and Stat1–knockout mouse embryo fibroblast (MEF) cell lines were used (34). Cells were maintained in DMEM supplemented with 10% FBS and 1% penicillin and streptomycin. Cells were plated at 60-70% density and maintained in full medium for a day. Cells were incubated in serum-free medium for another 24 hours. Cells were treated with TNF-*α* (20 ng/ml) or IFN-*β* (1000U/ml) for the indicated time. Cytokines were purchased from PBL Assay Science (Nutley, NJ). RAW 264.7 cells were treated with PolyIC (10µg/ml)), LPS (10ng/ml) or IFN-*β* (1000U/ml). PolyIC and LPS were purchased from InvivoGen (San Diego, CA) and used according to the manufacturer’s instructions.

### mRNA Expression and Western blot Analysis

RNA was extracted from cell pellets using the Qiagen RNeasy kit (Valencia, CA). Reverse transcriptase-polymerase chain reactions (RT-PCR) were performed using Ambion Retroscript (Austin, TX), according to the manufacturer’s protocol. Primer sequences for Egr-1, Irf1, and Gapdh were obtained from the molecular reagents section of Mouse Genome Informatics (MGI). Human Egr-1 and *β*-Actin primer sequences were previously described (22). PCR products were resolved on a 1% agarose gel containing ethidium bromide and visualized with U.V. Light and images were captured on a digital system. Image files were processed with ImageJ (NIH) software. Specific gene expression was normalized to GAPDH or *β*-Actin and fold changes in the treated samples were calculated with respect to controls. MLE-Kd cell extracts were prepared and proteins were separated by electrophoresis using 8%–10% SDS-PAGE gels. Proteins in the gel were electrophoretically transferred to polyvinylidene difluoride membranes (Bio-Rad, CA) and subjected to immunoblotting with the antibodies for phosphorylated Erk-1/2 (Thr202/Tyr 204) or total Erk-1/2 from Cell Signaling Technology (Beverly, MA). Blots were visualized by enhanced chemiluminescence western detection system (Pierce, IL).

## 3 Results and Discussion

### 3.1 Regulation of Egr-1 by Interferons and Toll-Like Receptor (TLR) ligands

The role of phosphorylation cascade and activation of the Jak-Stat pathway in IFN-*α*/*β* signaling is well established (1,2). In addition, inducible transcription factors such as Irf1, Irf2, Irf7 also play an important role in sustaining and modulating the IFN-*α*/*β* signaling (4,5). Transcription factor profiling in the transcriptome provides novel insights into the functional organization and connectivity in signal transduction pathways in eukaryotic cells (35). A large number of microarray studies on interferon-regulated gene expression were limited by factors such as cell type, long delay after treatment (2-24 hours) or treatment in cell culture media with high serum. These studies may have missed earlier dynamic changes in transcription factors and growth-regulated genes. In order to identify novel inducible transcription factors regulated by Interferon *α*/*β*, gene expression datasets representing early time points (0.5-2 hours) were examined (27,29). A wide variety of transcription factors were induced by IFN-*α*/*β* in human peripheral blood mononuclear cells (PBMC) and in Huh7 hepatoma cells (Figures 1A and 1B). Several metallothionein (MT) isoforms were induced in the early transcription response by IFN-*α*/*β* in Huh7 liver cells. Metals and inflammatory stimuli are the major inducers of metallothionein in the lung and liver. Early growth response-1 (Egr-1) was rapidly induced by IFN-*α*/*β* in both cell types and selected for further studies. RNA expression studies in mouse lung epithelial (MLE-Kd) and human fibrosarcoma (2fTGH) cell lines confirmed that IFN-*α*/*β* induced Egr-1 (Figure 2A and 2B). These results were consistent with previous studies in human fibroblasts (15). TNF-*α* induction of Egr-1 was much higher than IFN *α*/*β* in 2fTGH cells (Figure 2B). Interestingly, TNF-*α* but not IFN-*α* induced Egr-1 mRNA in Hela cells suggesting differential pathway regulation (15). Furthermore, IFN-λ induction of Egr-1 was higher than with IFN-*α*/*β* in Huh7 cells (Figure 2C). TLR recognition of viral and bacterial components such as double-stranded RNA (Poly IC) and LPS activate diverse signal transduction pathways, stimulate the production of type I Interferons and activate a large number of genes including interferon-stimulated gene expression (36). TLR4 ligand (LPS) and TLR3 ligand (Poly IC) induced Egr-1 mRNA in RAW264.7 mouse macrophage cells. Interestingly, LPS treatment resulted in the rapid and enhanced activation compared with Poly IC or IFN-*β* treatment in RAW 264.7 cells (Figure 3A). Time-course experiments in bone marrow derived-macrophages (BMDM) revealed distinct Egr-1 gene expression profiles for each treatment (Figures 3B-3D). These studies revealed that cytokines and TLR ligands use multiple and distinct signal transduction pathways to induce Egr-1 gene expression. Factors such as diversity, intensity and duration of the signal transduction pathways might account for the variation in Egr-1 expression levels in response to different cytokines and TLR ligands.

**Figure 1.**
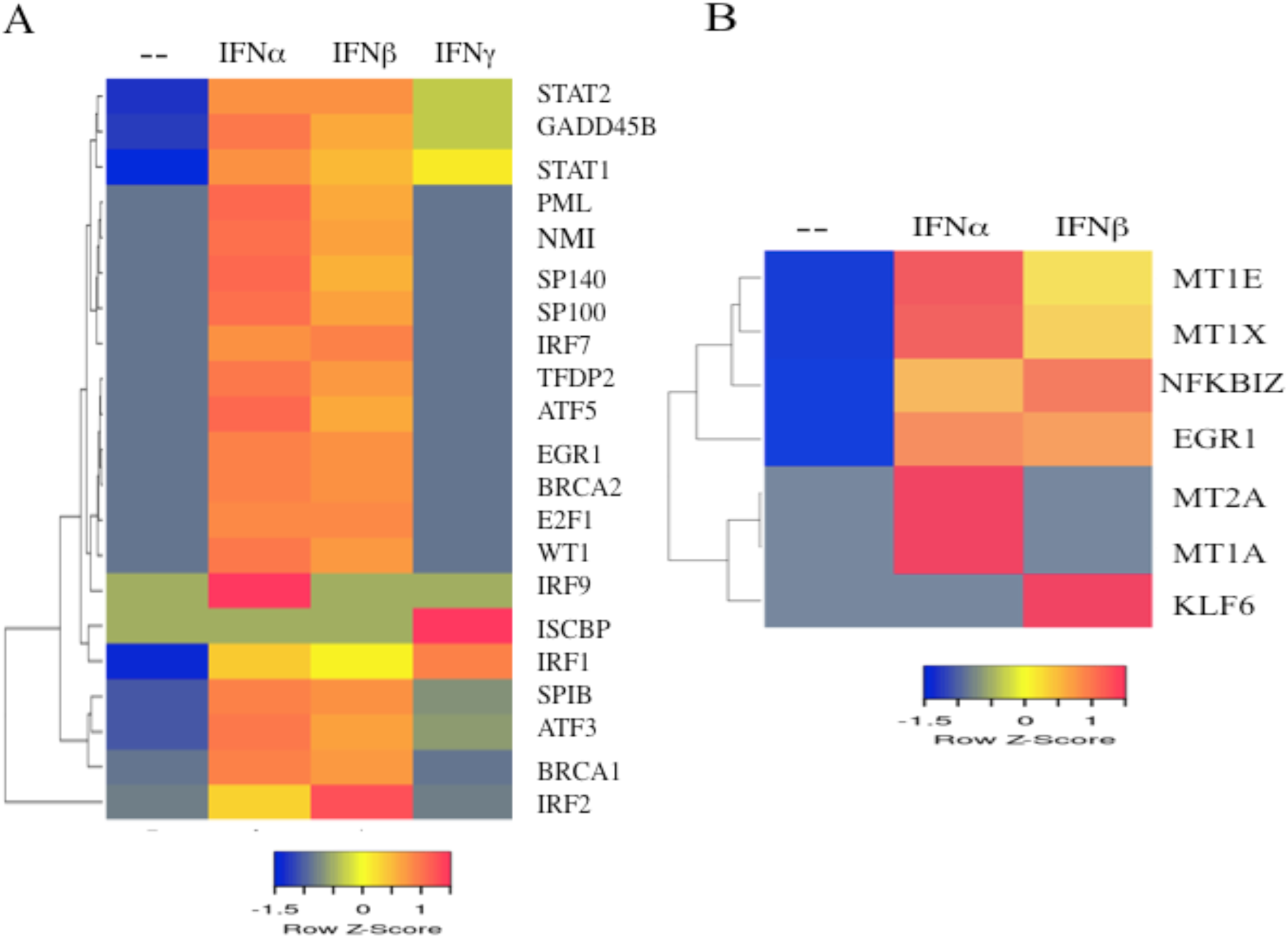
Regulation of transcription factor (TF) mRNA levels in human peripheral blood mononuclear cells (PBMC) and human hepatoma (Huh7) cells by interferon treatment (A) Transcription factor expression profiles were retrieved from microarray datasets of human PBMC treated with IFN-*α*, IFN-*β* or IFN-γ for 0.5-2 hours (B) Transcription factor expression in Huh7 cells treated with IFN-*α* or IFN-*β* for 0.5 hours were retrieved from the Microarray datasets using the GEO R2R analysis. Cluster analysis of gene expression was performed using the heatmapper software.

**Figure 2.**
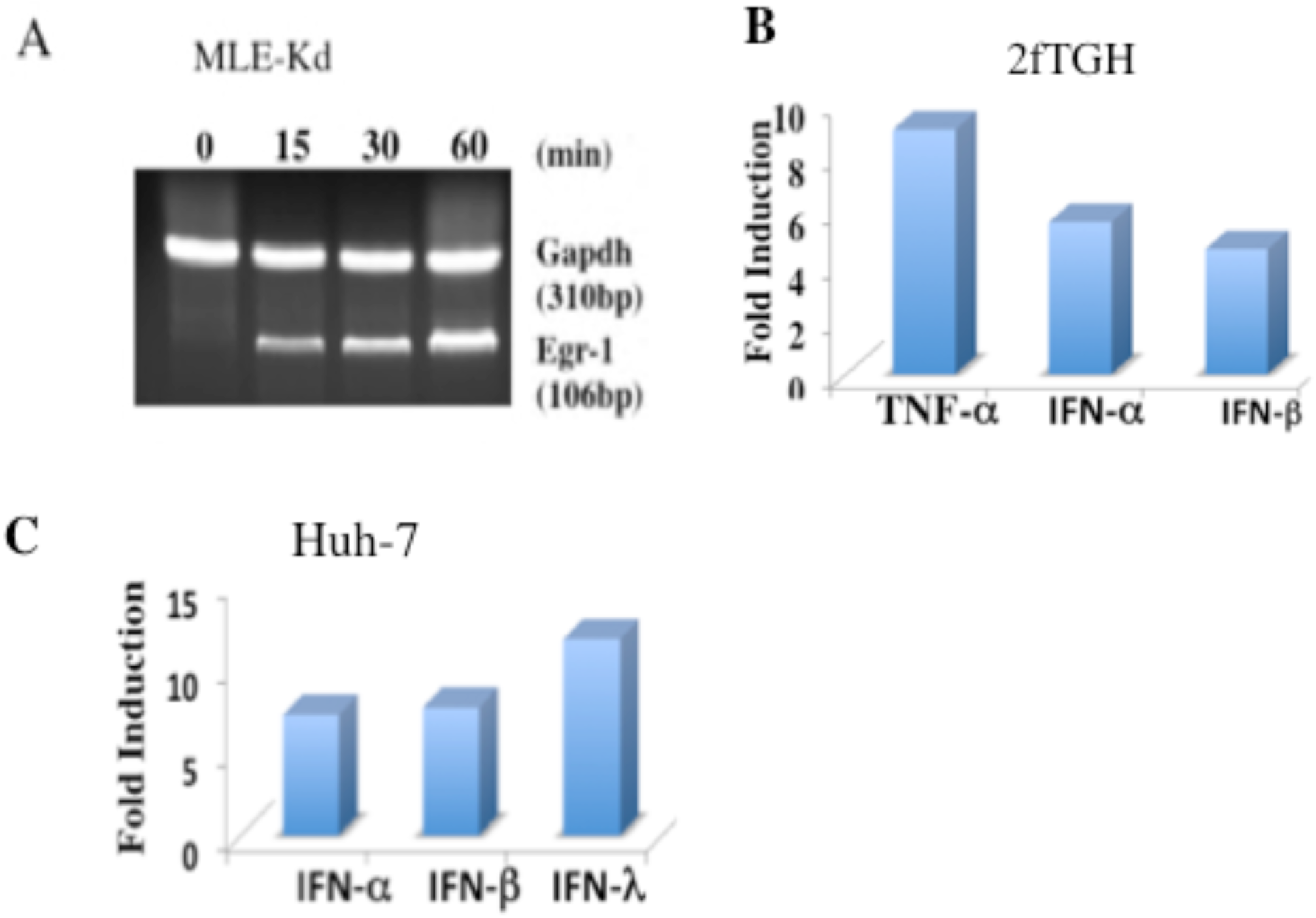
Regulation of Egr-1 mRNA levels by Interferon-Beta (IFN-*β*) in mouse and human cells. (A) Mouse MLE-Kd cells were treated with IFN-*β* (1000U/ml) and RNA was prepared from cells at 0, 0.25, 0.5, and 1 hour after treatment. RNA levels of Egr-1 and GAPDH were determined by RT-PCR analysis. (B) Human 2fTGH were treated with TNF-*α* (20 ng/ml), IFN-*α* (1000U/ml) or IFN-*β* (1000U/ml) for 2 hours. Egr-1 and *β*-Actin levels were determined by RT-PCR in control and treatment and expressed as fold induction. (C) Regulation of Egr-1 in Huh7 cells in response to IFN-*α*, IFN-*β*, and | FN-λ. Fold change values of Egr-1 were retrieved from Geo Datasets using the GEOR2R analysis.

**Figure 3.**
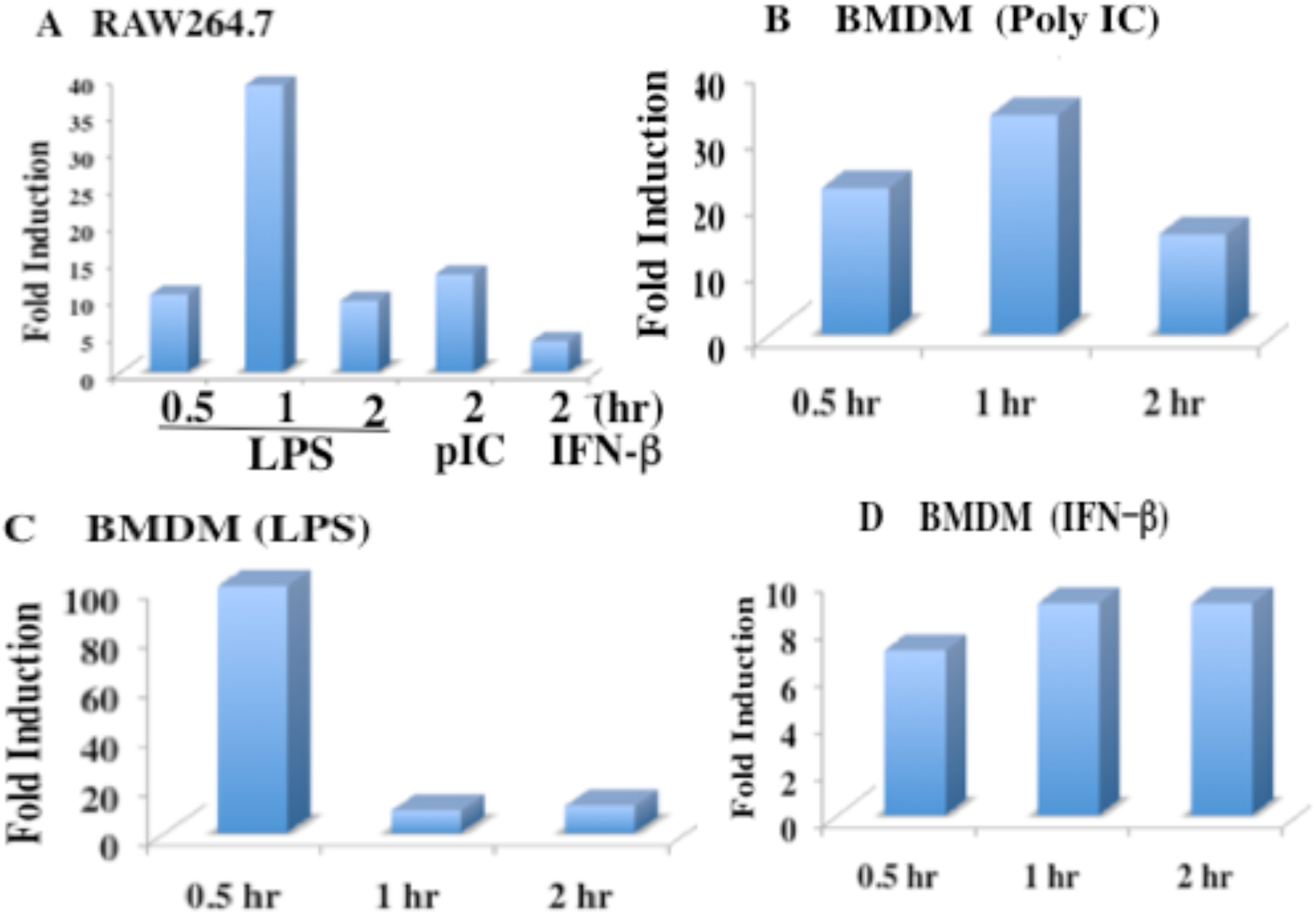
Regulation of Egr-1 mRNA levels in mouse macrophages by TLR ligands and IFN-*β*. (A) Mouse macrophage (RAW264.7) cells were treated with LPS for 0.5-2 hours or with polyIC or IFN-*β* for 2 hours. Egr-1 and GAPDH levels were determined by RT-PCR analysis in control and treatment and expressed as fold induction. (B-D) Egr-1 mRNA levels in BMDM cells treated with polyIC, LPS, or IFN-*β* for 0.5 to 2 hours. Egr-1-fold induction data were taken from reference 28.

### 3.2 Regulation of Egr-1 by IFN-*β* was independent of Stat1 or ISGF-3 components and mediated by the activation of Erk-1/2 pathway

IFN-*α*/*β* activates transcription factor complexes including Stat1 homodimer, Stat1-Stat2 heterodimer, and Stat1-Stat2-Irf9 trimeric complex (ISGF-3) that are involved in gene regulation (1,2). However, IFN-*γ*-mediated induction of Egr-1 was Stat1-independent in mouse embryonic fibroblasts (34). Studies in wild-type and Stat1-KO mouse fibroblasts showed that IFN-*α*/*β* induction of Egr-1 was independent of Stat1 (Figure 4A). In contrast, Irf1 induction by IFN-*α*/*β* was strictly Stat1-dependent. Furthermore, Egr-1 induction was 3-fold higher in Stat1-mull cells compared with wild-type cells. This enhanced expression of Egr-1was also observed in Stat1 KO fibroblasts in response to IFN-*γ* (34). Studies in mutant cells lacking Stat1 (U3A) Stat2 (U6A) and Irf9 (U2A) demonstrated that IFN-*β* induction of Egr-1 was independent of ISGF-3 components (Figure 4B). TNF–*α* and TLR ligands activate several Mitogen-activated protein kinase (MAPK) pathways such as Extracellular signal-regulated kinase (ERK), Jun N-terminal kinase (JNK) and p38 (37). Activation of the Erk-1/2 pathway leading to the phosphorylation of transcription factors such as Elk1 and Srf1 was implicated in the rapid induction of Egr-1 in response to multiple stimuli (26,38). Western blot analysis revealed that IFN-*β* rapidly activated Erk-1/2 within 30 minutes in serum-starved MLE-Kd cells (Figure 5A). Furthermore, pretreatment with Mek inhibitor (U0126) inhibited the induction of Egr-1 mRNA (Figure 5B). Pretreatment with an equal volume of DMSO (vehicle) has no significant effect on Egr-1induction. Activation of Jak1 and MAPK-interacting kinase (Mnk1) were required for the activation of Erk-1/2 in IFN-*β* signaling (39,40). There are technical complications in the analysis of Erk-1/2 phosphorylation and activation in cell lines including constitutive activation in transformed cells as well as cell culture conditions such as serum and nutrients (39-41). Erk-1/2 activates transcription factors such as Elk1 and Srf1 that bind to Ets and serum response elements (SRE) respectively, and mediate transcriptional activation of Egr-1 in response to serum and LPS (26,38). Mouse Egr-1 gene promoter contains multiple SRE elements as well as Activator protein (AP1), Cyclic AMP response (CRE), and Ets elements upstream of the transcription start site (Figure 5C). Activation of Srf1 and Elk1 by phosphorylation and binding of Srf1 and Elk1 to the Egr-1 gene promoter in response to IFN-*β* in MLE-Kd cells remains to be established. Expression of Srf1 enhanced the mRNA levels of several interferon-stimulated genes in response to Interferon-*α*/*β* in mouse macrophages (42). Detailed promoter analysis revealed that multiple distal and proximal cis-elements were involved in Egr-1 induction (43).

**Figure 4.**
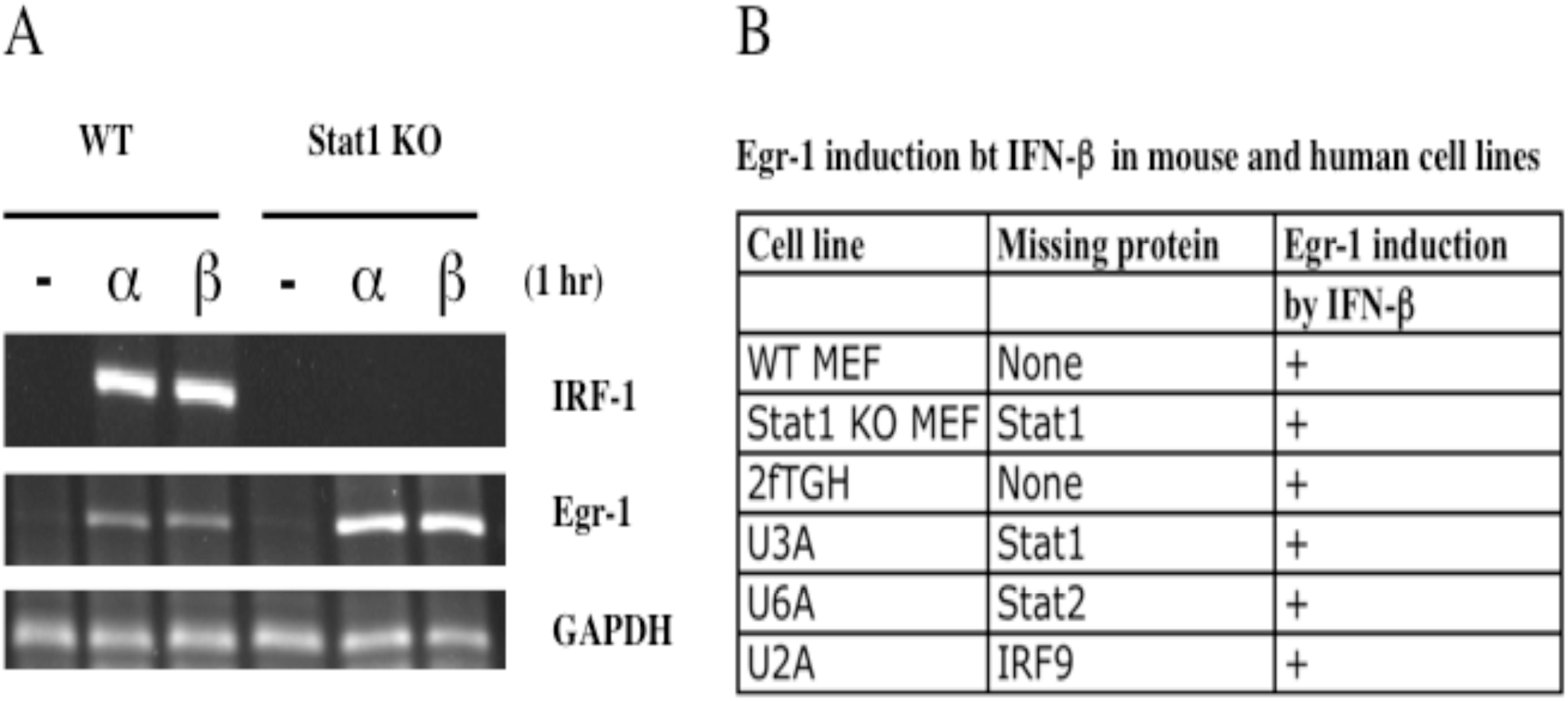
Regulation of Egr-1 mRNA levels by IFN-*β* in mouse and human mutant cell lines. (A) Wild-type and Stat1 KO MEF cells were left untreated or treated with IFN-*α* or IFN-*β* (1000U/ml) for one hour. RNA levels of Irf1, Egr-1, and GAPDH were determined by RT-PCR analysis (B) Summary of Egr-1 mRNA regulation by IFN-*β* in wild-type and Stat1 KO MEF cells and in human wild-type (2FTGH), Stat1-null (U3A), Stat2-null (U6A), and IRF9-null (U2A) fibrosarcoma cell lines.

**Figure 5.**
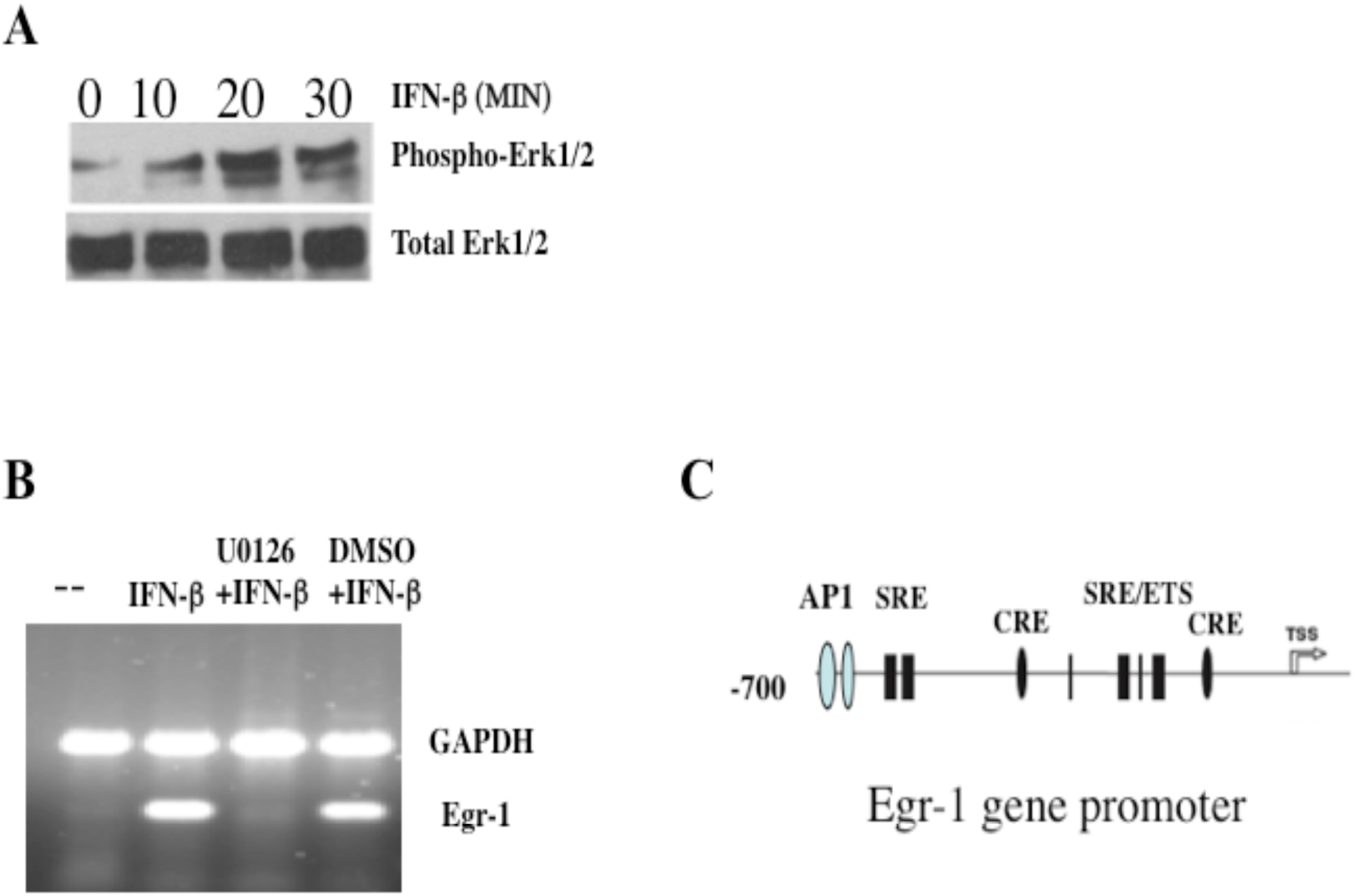
Activation of extracellular signal-regulated kinases (Erk1/2) by IFN-*β* in MLE-Kd cells (A) MLE-Kd cells were treated with IFN-*β* for 0, 10, 20 or 30 min. Cell extracts were prepared and the protein levels of phosphorylated Erk-1/2 and total Erk-1/2 were determined by western blot analysis. (B) MLE-Kd cells were left untreated or pre-treated with 40 µM of Mek-1 inhibitor U0126 or with an equal volume of DMSO for 2hours. Cells were stimulated with IFN-*β* alone or after pretreated with DMSO or U0126 for an additional 60 min. Total RNA was isolated and Egr-1 and GAPDH mRNA expression levels were determined by RT-PCR analysis. (C) Schematic representation of the mouse Egr-1 gene promoter. Location of the AP1, CRE, SRE, and Ets elements were shown. TSS represents the transcription start site. The map is not to scale.

### 3.3 Regulation of genes by Interferon-*α*/*β* and TNF-*α* in BMDM and PBMC

The generation of an inflammatory response is a complex process involving multiple cytokines acting in parallel and in concert in innate and adaptive immunity (36,44). Influenza virus-infected dendritic cells and macrophages, which reside in close proximity to lung epithelium, can produce significant amounts of TNF–*α* and type I IFN in response to virus infection (45). IFN-*α*/*β* is involved in signaling cross-talk with TNF-*α* that enhance or dampen the severity of inflammatory response (9,10). Interaction between Interferon–*α*/*β* and TNF-*α* signaling was reported in autoimmune diseases. (12,13). A select list of genes was induced by both TNF-*α* and Interferon-a/*β* in PBMC and BMDM gene expression datasets (Figure 6). TNF-*α* induction of these genes was much higher than IFN *α*/*β* in the mouse BMDM cells. Interestingly, Egr-1 was induced by both cytokines in PBMC and BMDM. These genes are involved in transcriptional regulation and integrating signal transduction pathways that are likely to play an important role in the cytokine storm and shift to hyperinflammatory gene expression in response to coronavirus infections (30,65).

**Figure 6.**
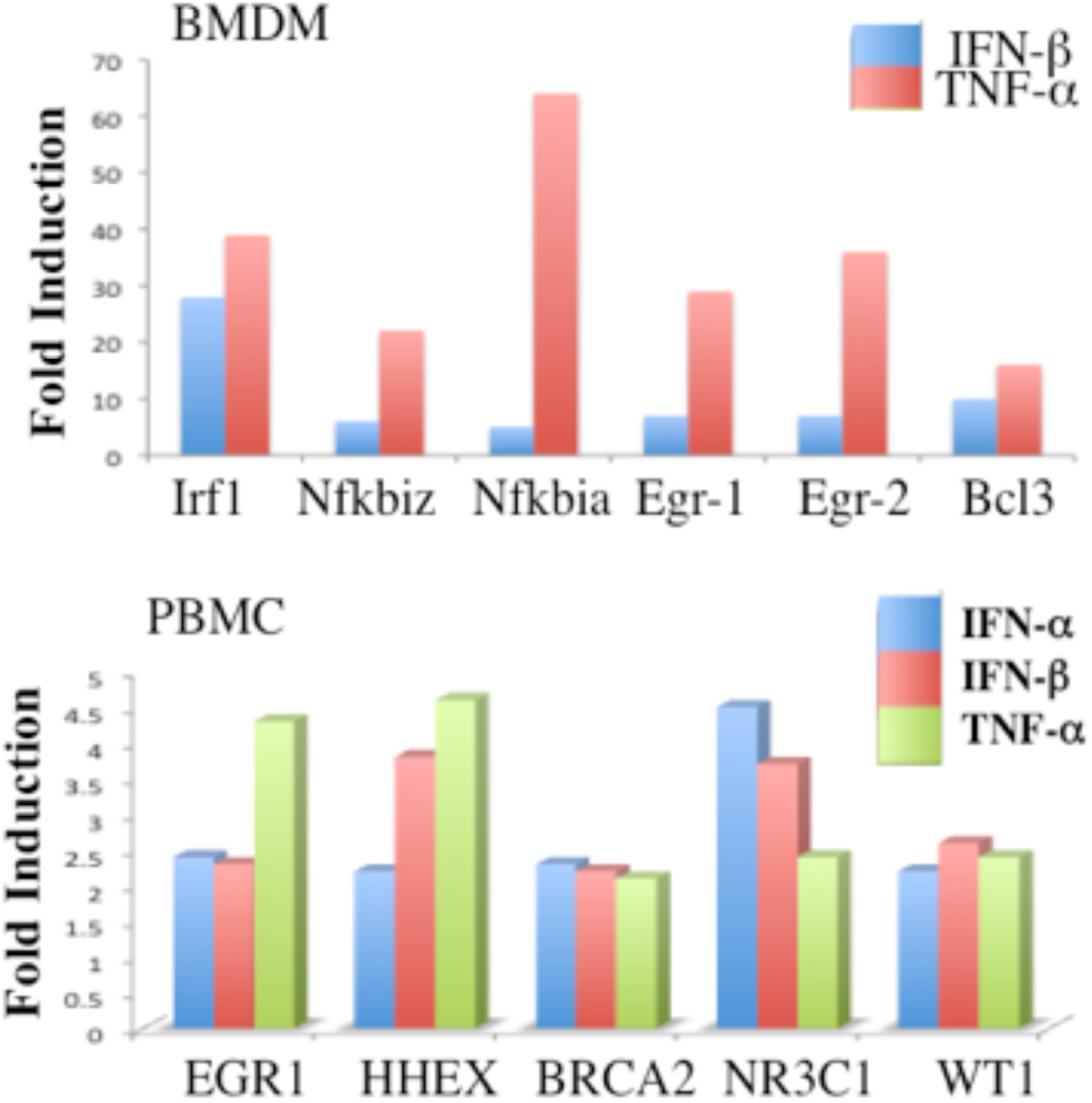
Selected list of genes involved in transcriptional regulation and signaling pathways regulated by Interferon-*α*/*β* and TNF-*α* in human PBMC and mouse BMDM cells. Gene expression profiles in PBMC (0.5-2 hours) and BMDM (0.5-2 hours) were retrieved from published array datasets.

### 3.3 Regulation of Egr-1 target genes in PBMC and BMDM by IFN-a/*β*

Egr-1 is a zinc finger transcription factor that binds to a GC-rich sequence in the promoter regions of target genes. A shortlist of Egr-1 regulated genes was generated from the TRUUST transcription factor database and published studies in different cell types and under different stimuli (20-22,46). Egr-1 regulates genes involved in cell growth (Egr2, Atf3, Pdgfrb, Cdkna1, Ccnd1, Tnfsf10, and chemokines involved in inflammation such as (Cxcl10, Cxcl2, Ccl2, Ccl3). It is important to note that many of the Egr-1 target genes were also known targets of NF-KB, AP1, and Stat1 in cytokine signaling (46). Furthermore, Egr-1 interacts with several other transcription factors such as Ets1, Ets2, Elk1 that are common in cytokine responsive genes (46). Previous studies have shown that transcription factors of innate and adaptive immunity show functional connectivity involving shared interacting protein partners and target genes. (47). Cluster analysis revealed that Egr-1 induction was correlated with the expression of Egr-1 target genes by IFN-*α*/*β* in PBMC and in BMDM cells (Figure 7).

**Figure 7.**
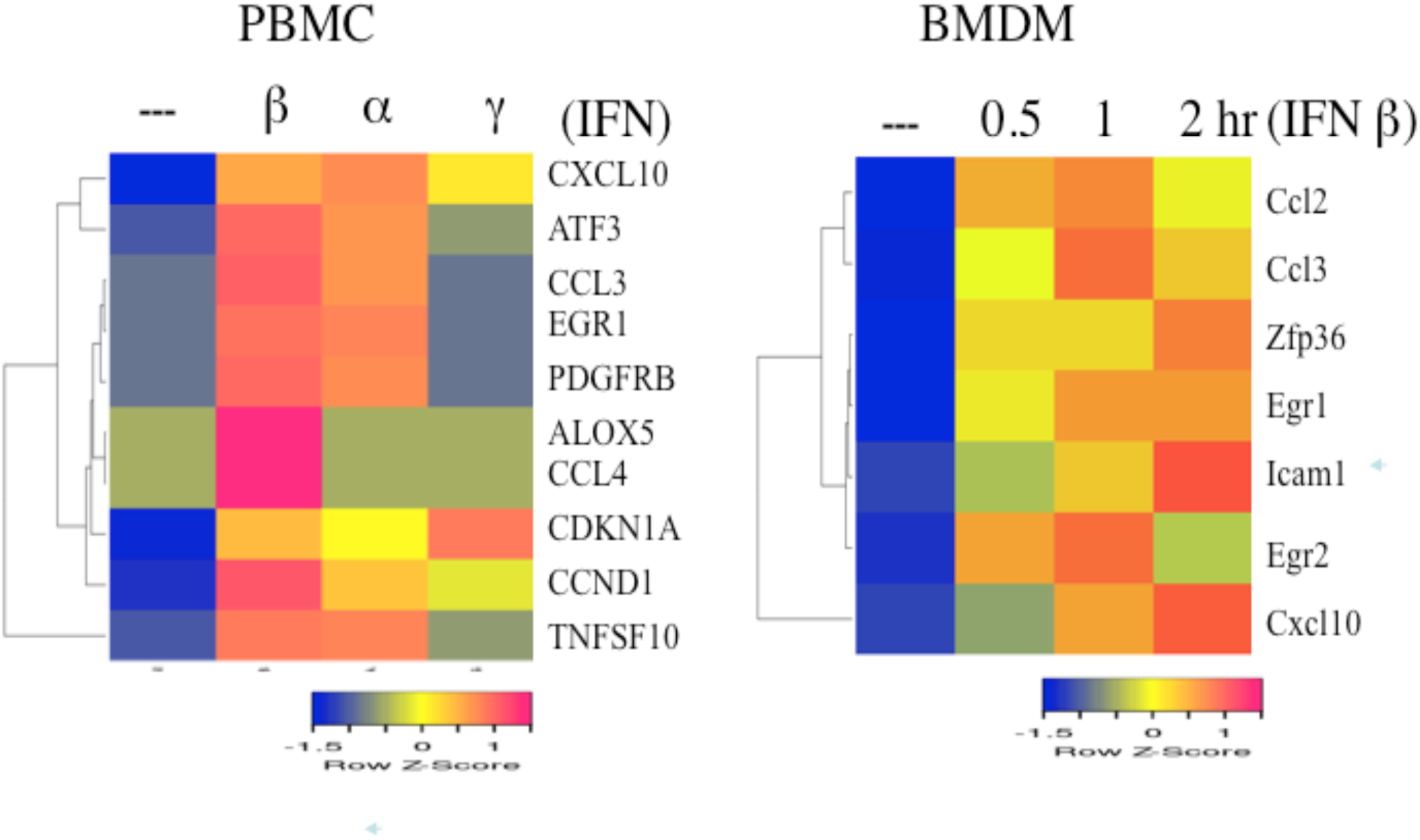
Regulation of mRNA levels of Egr-1 target genes by interferon treatment in human PBMC and mouse BMDM cells (A) Egr-1 target gene expression in human PBMC treated with IFN-*α*, IFN-*β* or IFN-*γ* for 0.5-2 hours (B) Egr-1 target gene expression in mouse BMDM cells treated with IFN-*β* for 0.5-2 hours. Gene expression profiles were retrieved from published array datasets. Cluster analysis was performed with the heatmapper software.

### 3.4 Regulation of Egr-1 By Severe Acute Respiratory Syndrome Coronaviruses (SARS-CoV) and other Respiratory Viruses

Severe acute respiratory syndrome coronaviruses (SARS-CoV) and influenza viruses are major respiratory pathogens in humans with seasonal epidemics and potential pandemic threats (48,49). Respiratory virus infection results in lung injury, a significant component of which was mediated by the host immune response (50). The highly pathogenic 1918 Influenza virus-induced gene expression signature in mouse lungs characterized by enhanced expression of cytokines such as TNF-*α*, IFN-*γ*, and IL-6 resulting in a dramatic increase in chemokines and inflammatory cell influx consisting of neutrophils, macrophages, and lymphocytes (51,52). Gene expression profiling in the lungs of mice infected with 1918 influenza HINI showed enhanced expression of Egr-1 and its downstream target genes Cxcl2 and Ccl2 involved in neutrophil and macrophage inflammatory influx, respectively (51). Activation of multiple signal transduction pathways leading to the activation of transcription factors such as AP1 and NF-kB in response to influenza infection were reported (53,54). Influenza virus propagation was impaired by inhibition of the Mek/Erk pathway and lung-specific expression of active Raf kinase results in increased mortality of influenza virus-infected mice (55,56). Enhanced cytokine and chemokine expression were also reported in longitudinal studies of severe COVID -19 patients (57).

Gene expression profiling studies of human lung epithelial cell lines such as Calu-3, A549, and NHBE1 in response to respiratory viruses were reported recently (30,31). Interrogation of Microarray data sets revealed that Egr-1 was induced by SARS-CoV-1 and SARS CoV-2 infection but not by mock-infection in Calu-3 cells. This induction was observed at 24 hours after infection. Egr-1 regulated gene TNFSF10 (TRAIL) was also induced by SARS-CoV-1 and SARS CoV-2 infection (Figure 8). TRAIL is a widely expressed member of the TNF family involved in critical biological functions including apoptosis and tumor suppressor activities (58). TRAIL activation was demonstrated to be part of the effector mechanisms by which influenza-specific T cells protect the host against virus infection (54,59). Furthermore, Egr-1 target genes including transcription factors (ATF3) and chemokines (CXCL10) were also induced by SARS-CoV-1 and SARS CoV-2 infection of Calu-3 cells (Figure 9). In contrast, SARS-CoV-2 infection of A549 failed to induce Egr-1 expression (data not shown). Recent studies have shown that Angiotensin-converting enzyme 2 (ACE2) functions as ectopeptidase and virus entry receptor and facilitates entry of SARS-CoV1 and SARS-CoV-2 in lung epithelial cells (60). Expression of ACE2 in A549 cells rescued SARS-CoV-2 -mediated Egr-1 induction and the expression of Egr-1 target genes such as ATF3 (Figure 10). Furthermore, expression of CXCL10, CXCL2, CCL3, CCL4 chemokines were dramatically increased after SARS-CoV-2 infection of A549 cells expressing ACE2 (Figure 11 and data not shown). Importantly, the expression of ACE2 alone in A549 cells did not induce Egr-1 mRNA levels. Differential regulation of SARS-CoV-2 in Calu-3 and A549 suggest that these cell lines may differ in the expression levels of host factor (ACE2) required for SARS-CoV-2 entry. Pandemic 1918 H1N1 influenza virus infection induced expression of Egr-1 and Egr-1 dependent chemokines in mouse lungs (Figure 12A). In contrast, influenza A virus (IAV) suppressed Egr-1 expression in NHBE bronchial epithelial cells that is dependent on viral non-structural protein or NS1 (Figure 12B). These studies suggest that multiple factors such as influenza strain, virus-encoded factors (NS1), cell type regulate Egr-1 expression. Respiratory syncytial virus (RSV) or parainfluenza virus (HPIV3) infection also induced Egr-1 mRNA in A549 cells, 24 hours post-infection (Figure 12C). These studies suggest that Egr-1 expression is a common host response to many respiratory viruses

**Figure 8.**
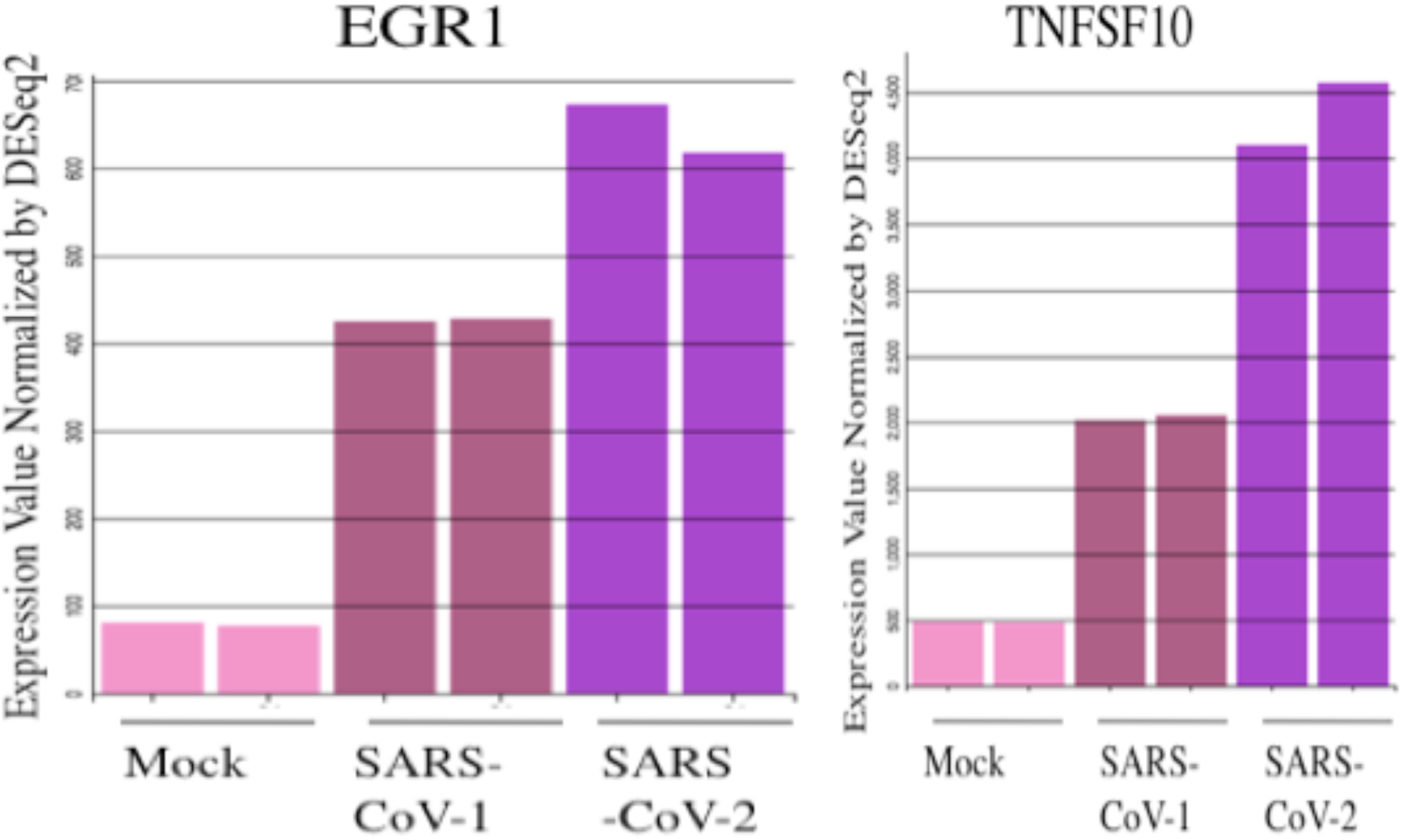
Regulation of Egr-1 and TNFSF10 mRNA levels by Coronaviruses in Calu-3 cells. Calu-3 cells were mock-infected or infected with the virus (SARS-CoV-1 or SARS-CoV-2) for 24 hours. Data from two samples for each condition were shown. Expression levels were normalized by DEseq2.

**FIgure 9.**
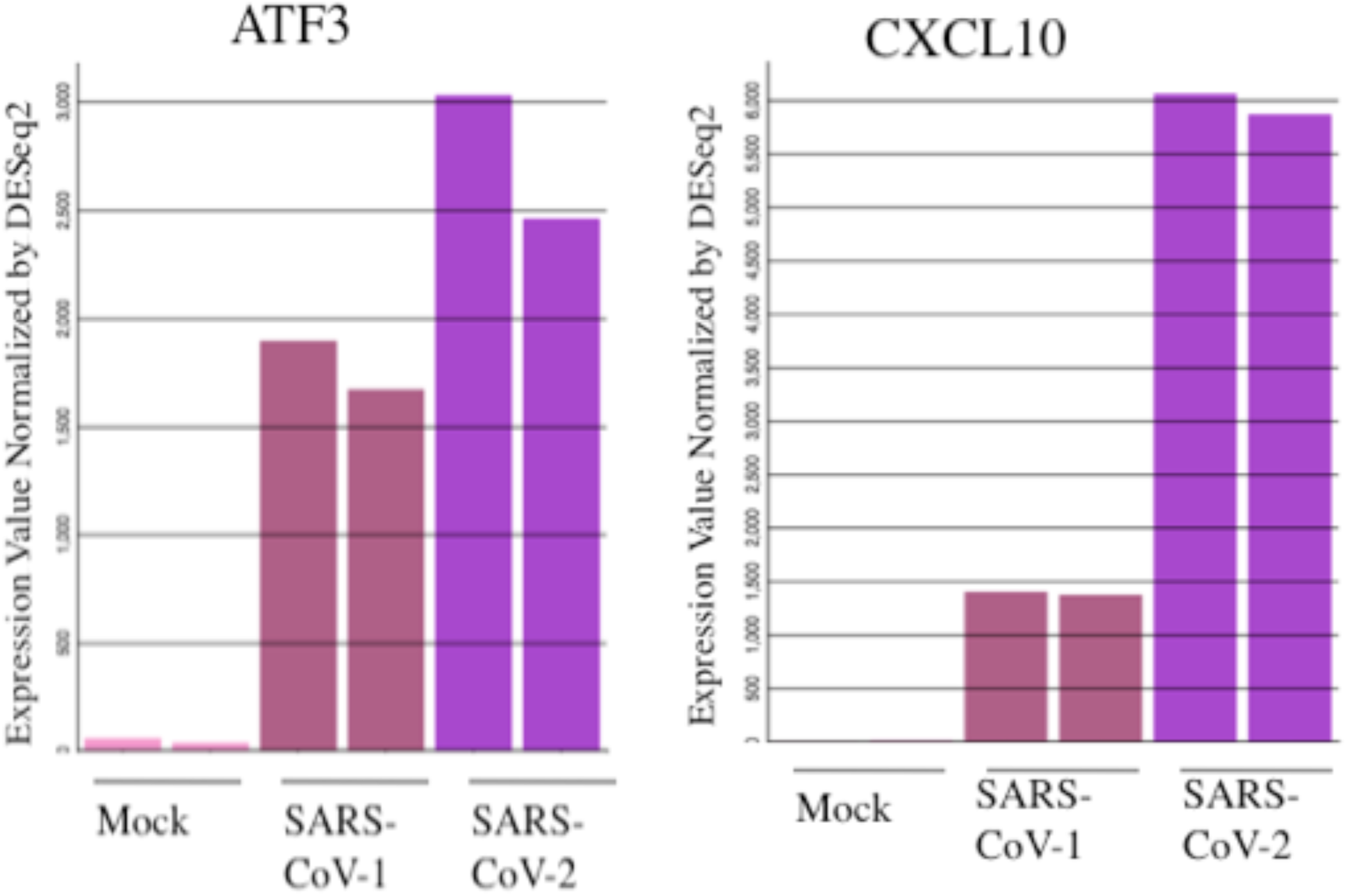
Regulation of Egr-1 inducible genes ATF3 and CXCL10 mRNA levels by Coronaviruses in Calu-3 cells. Calu-3 cells were mock-infected or infected with the virus (SARS-CoV-1 or SARS-CoV-2) for 24 hours. Data from two samples for each condition were shown. Expression levels were normalized by DEseq2.

**Figure 10.**
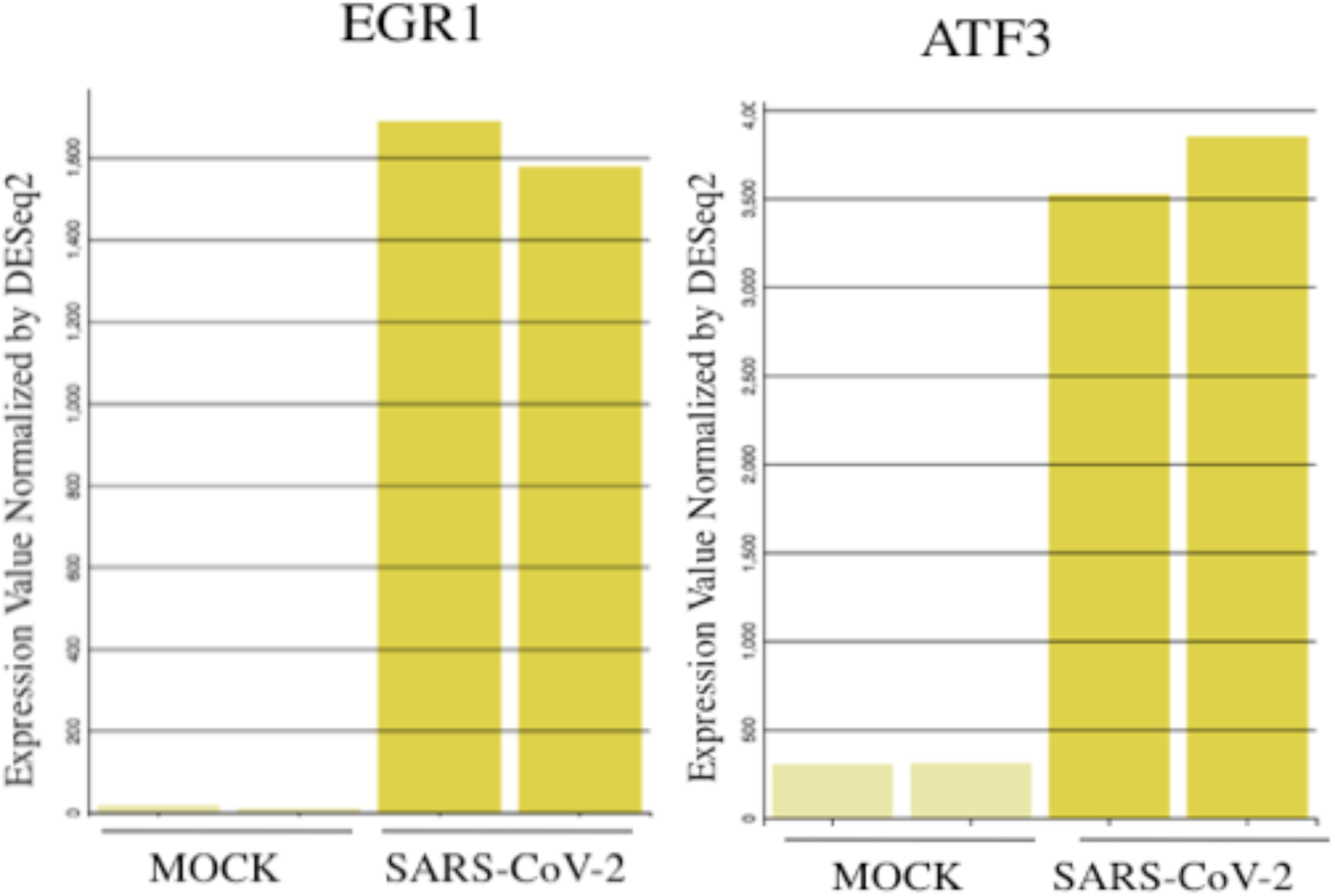
Regulation of Egr-1 and ATF3 transcription factor mRNA levels by SARS-CoV-2 in A549 cells expressing ACE2 receptors. Cells expressing ACE2 receptors were mock-infected or infected with SARS-CoV-2 virus for 24 hours. Egr-1 and ATF3 mRNA expression levels were normalized by DEseq2. Data from two samples for each condition were shown.

**Figure 11.**
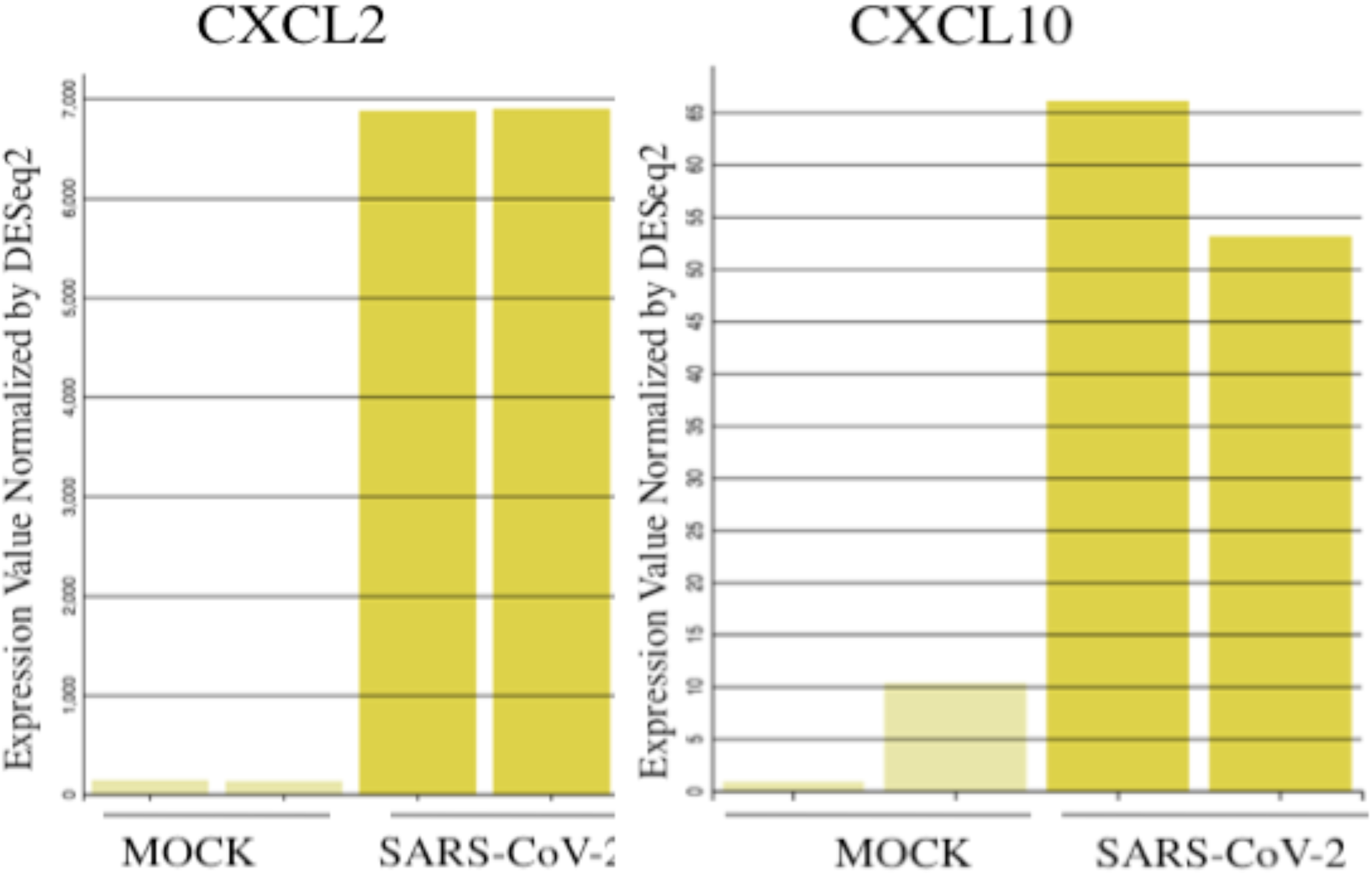
Regulation of Egr-1 inducible chemokines CXCL10 and CXCL2 mRNA levels by SARS-CoV-2 in A549 cells expressing ACE2 receptors. Cells expressing ACE2 receptors were mock-infected or infected with SARS-CoV-2 virus for 24 hours. CXCL10 and CXCL2 mRNA expression levels were normalized by DEseq2. Data from two samples for each condition were shown.

**Figure 12.**
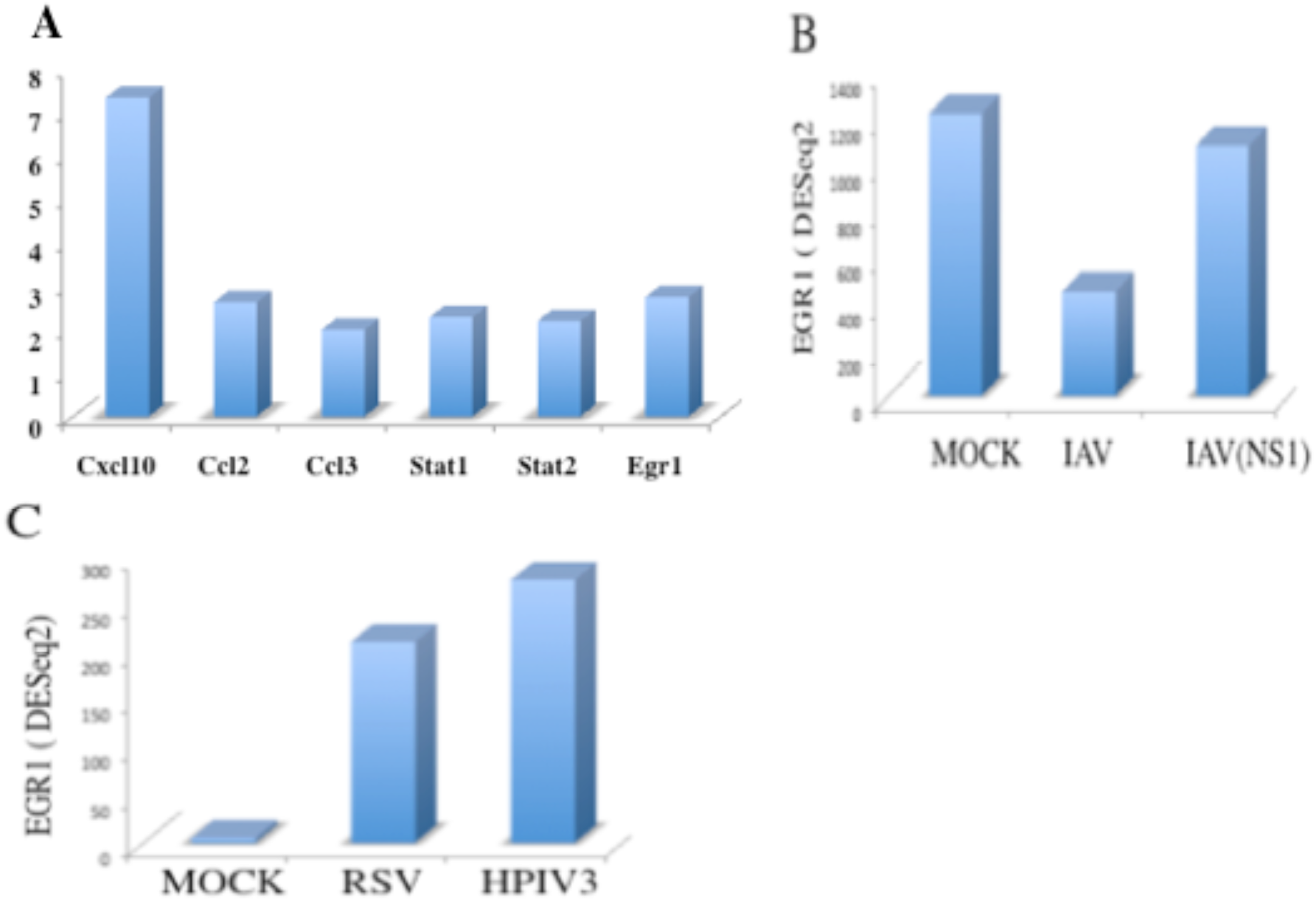
Regulation of Egr-1 mRNA levels by influenza viruses, respiratory syncytial virus (RSV) and parainfluenza virus (HPIV3). (A) Fold change in RNA expression levels of chemokines and transcription factors in the lung tissue of mice, three days after infection with reconstructed 1918 H1N1 influenza virus versus mock-infected controls. (B) Human NHBE1 bronchial epithelial cells were mock-infected or infected with influenza virus (IAV) or influenza virus lacking NS1 (IAVNS1) for 12 hours. (C) Human A549 lung cells were mock-infected or infected with RSV or PIV for 24 hrs. Egr-1 mRNA levels were normalized by DESeq2. Figure 13. Relative mRNA expression levels of Egr-1 and Cyclin D1 (CCND1) in the lung biopsies of healthy controls and COVID-19 patients. Healthy and COVID-19 samples were represented by black and red, respectively. RNA expression values were normalized by DEseq2. Data from two samples for each condition were shown.

### 3.5 Regulation of Egr-1 in COVID-19 patient lung biopsies and Peripheral Blood Mononuclear Cells

Gene expression profiling studies of a limited number of human lung biopsies and PBMC of healthy controls and COVID-19 patients were reported recently (30,31). Interrogation of the gene expression data in lung biopsies revealed that Egr-1 expression levels were significantly lower in COVID-19 patients compared with healthy controls. Expression of Egr-1 target genes related to cell growth such as CCND1, EGR2, and PDGFRB were also down-regulated (Figure 13 and data not shown). In contrast, expression of Egr-1 target genes involved in inflammatory responses such as CXCL10, CCL2, CCL3, CCL4, and TNFSF10 were dramatically enhanced in COVID-19 patients compared with healthy controls (Figure 14 and data not shown). Interestingly, STAT1 expression levels were significantly increased in COVID-19 patients compared with healthy controls (Figure 14). There are several limiting factors to interpreting the data. The lung is a complex tissue consisting of more than thirty cell types with differential contributions with respect to cell mass and gene expression. For example, type I and type II cells in mouse lungs constitute the major and minor cell types and express distinct cell markers (35). Without information on the expression levels in distinct cell types, it is difficult to correlate Egr-1 expression with target genes in the whole lung tissue. In support of this view, in a mouse model of CD8^+^ T cell -mediated lung injury, chemokine expression was dependent on Egr-1 activation in alveolar type II cells (63). Another intriguing possibility is that enhanced STAT1 expression compensates for decreased Egr-1 expression in some cell types in the lung. It is important to consider that in addition to changes in mRNA levels, the phosphorylation status of STAT1 and Egr-1 may play an important role in the transcriptional regulation of chemokine target genes. It is likely that differential expression in distinct cell types may account for the disparity of Egr-1 and its target gene expression in healthy controls and COVID-19 patients. Longitudinal studies in COVID-19 patients revealed that cytokines and chemokines were dramatically elevated in the lungs and correlated with the severity of the disease (57). Egr-1 mRNA levels were elevated and correlated with TNFSF10/TRAIL expression in the adult but not developing neutrophils of the peripheral blood mononuclear cells of COVID-19 patients compared with healthy controls (Figure 15). These results suggest the possibility of differential growth, survival, and apoptosis of mature versus developing neutrophils. Preliminary gene expression data in the lungs and PBMC of healthy and COVID-19 patients were limited by the number of datasets and should be interpreted with caution and needs further verification.

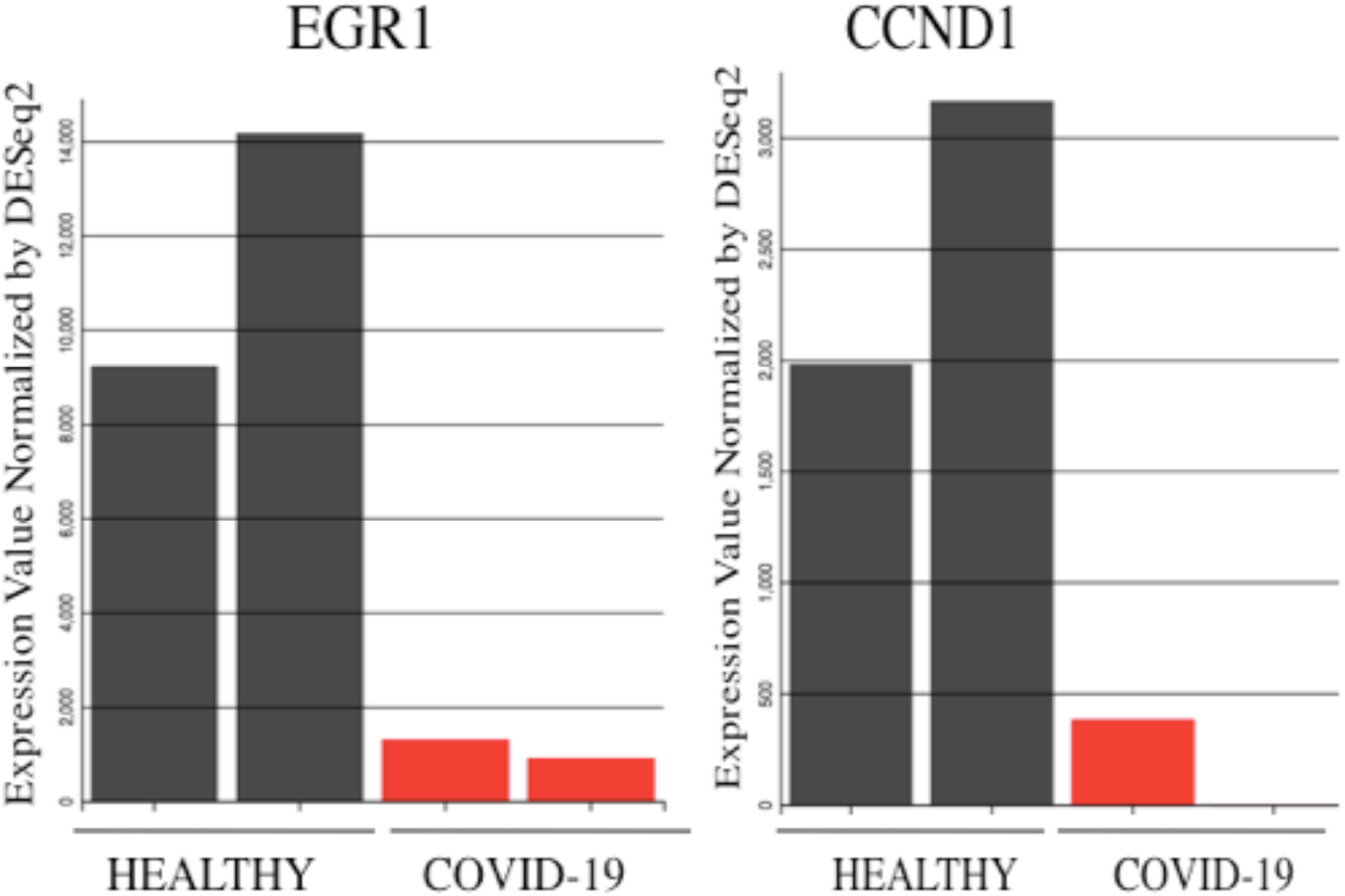

**Figure 14.**
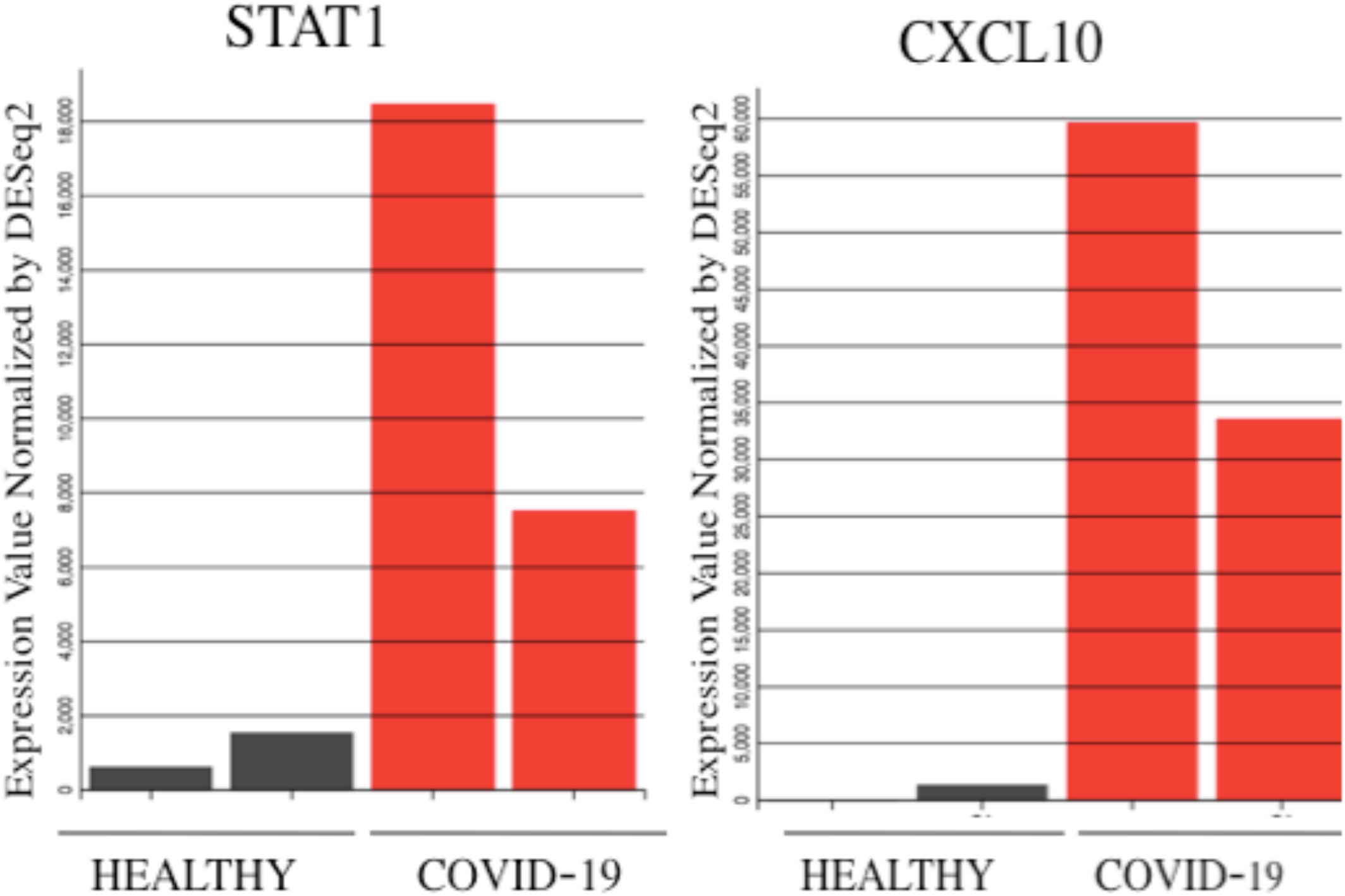
Relative mRNA expression levels of STAT1 and CXCL10 in lung biopsies of healthy controls and COVID-19 patients. Healthy and COVID-19 samples were represented by black and red, respectively. RNA expression values were normalized by DEseq2. Data from two samples for each condition were shown.

**Figure 15.**
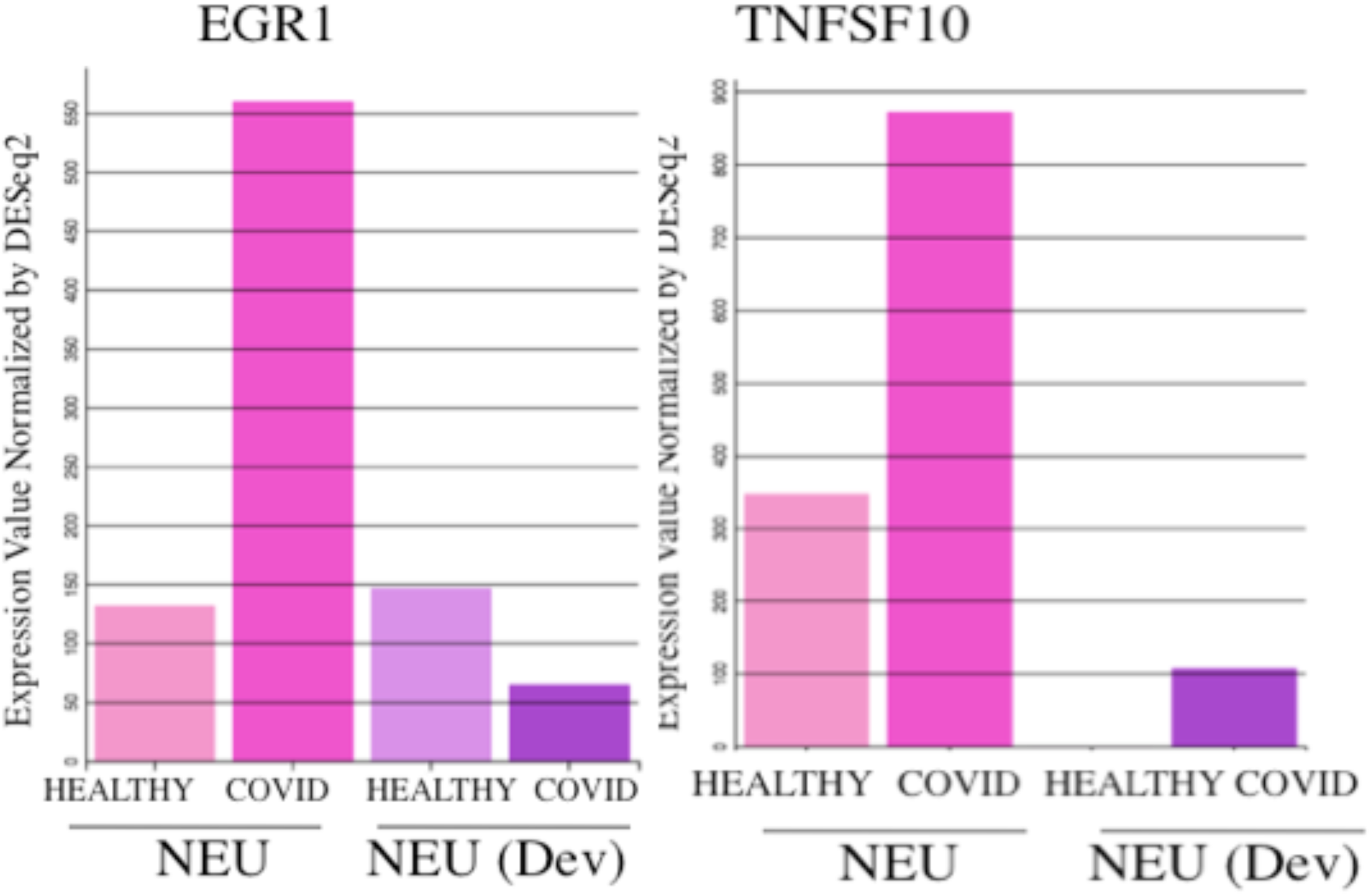
Regulation of Egr-1 and TNFSF10 mRNA levels in neutrophils of healthy controls and COVID-19 patients. Egr-1 and TNFSF10 RNA expression levels in mature Neutrophils (Neu) and developing Neutrophils (Neu Dev) of healthy controls and COVID-19 patients were shown. Egr-1 and TNFSF10 mRNA expression values were normalized by DEseq2.

## 4 Conclusion

Egr-1 has emerged as a key regulator of cell growth, reproduction, and response to tissue injury (16, 61-63). Egr-1 was rapidly induced by interferons and pro-inflammatory cytokines such as TNF-*α*, and IL-1*β* (15,63,64). Recent studies have demonstrated dramatic changes in type I interferon, TNF-*α*, and IL-1*β* production by immune cells and cytokine-mediated lung inflammation in COVID-19 patients (30,57,65,66). However, the role of Egr-1 in innate immunity and antiviral response to respiratory viruses in general and SARS-CoV-2, in particular, remains to be investigated. Transcriptional factor profiling in the transcriptome revealed that Egr-1was induced by IFN-*α*/*β*, TLR ligands, and TNF-*α* in human and mouse cells. Studies in mutant cell lines lacking Stat1, Stat2, and Irf9 revealed that IFN-*α*/*β* induction of Egr-1 was independent of ISGF-3 components and mediated by the activation of Erk-1/2 pathway. Furthermore, respiratory viruses such as SARS-CoV-2 induced Egr-1 and its target genes in several lung epithelial cell lines and in COVID-19 patients. Activation of Egr-1 by IFN-*α*/*β* and TNF-*α* and cross-talk between the pathways modulates signal transduction and inflammatory response in innate immunity.

## Acknowledgements

I would like to thank Dr. George Stark (Lerner Research Institute, Cleveland Clinic, OH) and Dr. Richard Enelow (Dartmouth-Hitchcock Medical Center, Lebanon, NH) for cell lines and research support.

